# Do Fossorial Water Voles have a Functional Vomeronasal Organ? A Histological and Immunohistochemical Study

**DOI:** 10.1101/2023.09.28.560035

**Authors:** Sara Ruiz-Rubio, Irene Ortiz-Leal, Mateo V. Torres, Aitor Somoano, Pablo Sanchez-Quinteiro

## Abstract

The fossorial water vole, *Arvicola scherman*, is a herbivorous rodent that cause significant agricultural damages. The application of cairomones and alarm pheromones emerges as a promising sustainable method to improve its integrated management. These chemical signals would induce stress responses that could interfere with the species regular reproductive cycles and induce aversive reactions, steering them away from farmlands and meadows. However, there is a paucity of information regarding the water vole vomeronasal system, both in its morphological foundations and its functionality, making it imperative to understand the same for the application of chemical communication in pest control.

This study fills the existing gaps in knowledge through a morphological and immunohistochemical analysis of the fossorial water vole vomeronasal organ. The study is primarily microscopic, employing two approaches: histological, using serial sections stained with various dyes (Hematoxylin-Eosin, PAS, Alcian Blue, Nissl), and immunohistochemical, applying various markers that provide morphofunctional and structural information. These procedures have confirmed the presence of a functional vomeronasal system in fossorial water voles, characterized by a high degree of differentiation and a significant expression of cellular markers indicative of active chemical communication in this species.

## INTRODUCTION

The fossorial water vole, *Arvicola scherman* (former fossorial form of *Arvicola terrestris* = *Arvicola amphibius*; for taxonomic considerations see Balmori-de la Puente et al. (2022) and references therein) is a cricetid rodent that lives in extensive burrow systems and preferentially consumes roots, bulbs and tubers of dicotyledons and some Poaceae (Airoldi 1976). This species inhabits grasslands and orchards in the main mountain massifs of Europe, including the northern Iberian Peninsula, the Alps, the mountains of central Europe and the Carpathians, and is considered one of the most important agricultural pests in these areas (Somoano 2020). Fossorial water voles show a high reproductive potential (Somoano et al. 2016; Somoano et al. 2017) and can reach high population densities of more than 1,000 individuals per hectare during population peaks (Giraudoux 1997), being able to spread unhindered in grassland-dominated landscapes (Somoano et al. 2022). This species may also pose a danger to human health because it can play an important role as reservoir of zoonotic pathogens such as *Leptospira* spp. (Giraudoux et al. 2009) or *Borrelia burgdorferi s.l.* (Espí et al. 2017), and parasites such as *Toxoplasma gondii* (Fuehrer et al. 2010) or *Echinococcus multilocularis* (Robardet et al. 2011). Cyclic vole species, such as fossorial water voles, have received much attention from rodent control organizations and their management requires effective, specific, environmentally benign, economically feasible and socially acceptable approaches (Jacob 2013).

Recognizing danger through chemical signals is of utmost importance for species survival (Lima and Bednekoff 1999). An avoidance response of water voles to the odors of the American mink, *Neovison vison*, has been observed in the wild (Brzeziński et al. 2019) and in outdoor enclosures (Barreto and Macdonald 1999; Nazarova et al. 2016). Since the fossorial water vole is primarily a subterranean species, chemical senses play a dominant role compared to physical senses like hearing and vision (Dennis et al. 2020). This detection is potentially mediated by kairomones, generally defined as substances producing a beneficial effect in a species other from the one producing them (Wood 1983). Kairomones are released naturally by predators and detected by their prey (Sbarbati and Osculati 2006), and differ from pheromones in that the latter act intraspecifically and influence behavior and reproductive functions (Wyatt 2003). However, alarm pheromones have been identified that alert conspecifics to the presence of predators (Stowe et al. 1995). A single substance can act as both a kairomone and an alarm pheromone, such as 2,4,5-trimethylthiazoline (TMT) which has been found in mouse feces (Brechbuhl et al. 2013) and fox urine (Rosen et al. 2015). The response to TMT manifests as elevated blood cortisol levels, systemic stress, and significant behavioral alterations, such as a change in habitat use (Dielenberg and McGregor 2001; Horii et al. 2010; Takahashi 2014). Another sulfur compound implicated in the aversive response to predators is 2-propylthietane, 3-methyl-1-butanethiol (Sievert and Laska 2016), and methyl-2-phenylethyl sulphide, found in urine of both male and female red foxes (McLean et al. 2021). These facts pave the way for the use chemical-based strategies that could be applied to improve integrated pest management in this species (Apfelbach et al. 2005; Papes et al. 2010).

The two fundamental chemical senses involved in chemical communication are the main olfactory system (MOS) (Salazar et al. 2019) and the vomeronasal system (VNS) (Halpern and Martínez-Marcos 2003; Torres et al. 2023b). The olfactory system is responsible for detecting olfactory chemical compounds involved in associative behaviors and learning (Firestein 2001; Menini 2010), and operates concurrently with the vomeronasal system (VNS), which is the olfactory pathway responsible for detecting pheromones (Liberles 2014), kairomones (Fortes-Marco et al. 2013), and molecules of the major histocompatibility complex (Leinders-Zufall et al. 2014). The olfactory molecules are recognized by receptors from the nasal conchae (Barrios et al. 2014b; Barrios et al. 2014b) and the information is then relayed to the main olfactory bulb (MOB) (Ennis and Holy 2015), which integrates and sends it to various brain areas (Price and Powell 1971; Brunjes et al. 2016). The VNS comprises a complex sensory structure called the vomeronasal organ (VNO), which is a paired structure located at the base of the nasal septum and is composed of a neurosensory epithelium lining a duct situated in a tubular formation communicating with the external environment (Salazar and Sanchez-Quinteiro 1998; Salazar et al. 2016). The transmission of the chemosensory information detected by the VNO is facilitated by unmyelinated nerve afferents whose terminations form the nerve layer of the accessory olfactory bulb (AOB) (Mori et al. 1987; Ortiz-Leal et al. 2022b). The AOB, in turn, projects to secondary centers, mainly to the vomeronasal amygdala (Scalia and Winans 1975; Mohrhardt et al. 2018).

Characterization of the VNS is a fundamental aspect of understanding the transmission of chemosensory information in fossorial water voles. However, to our knowledge, information on their anatomy and neurochemistry is lacking, apart from recent morphological information reported on odor-mediated sexual discrimination and attraction (Poissenot et al. 2023). Therefore, we conducted in this study an exhaustive morphological and immunohistochemical characterization of the *A. scherman* VNO to better understand its relevance in this species and its potential role in kairomone detection. Both macro- and microdissection methodologies were used along with general and specific histological staining techniques. These were enhanced by lectin-histochemical labeling and immunohistochemical staining. Results obtained here on the anatomy and neurochemistry of the fossorial water vole VNO were compared with those results obtained in other rodents such as rats (Salazar and Sánchez Quinteiro 1998), mice (Keller et al. 2022) and capybaras (Torres et al. 2020), or carnivores like minks (Salazar et al. 1998) and fox (Ortiz-Leal et al. 2020) to go deeper on its differentiation and functionality levels. This will provide the basis for future studies focused on assessing the physiological effects of kairomones on individuals, and ultimately explore practices aimed at containing the growth and spread of wild populations.

## METHODS

### 2.1 Specimen collection

A total of 10 specimens of *A. scherman* were captured (half of each sex) in grasslands located in the village of O Biduedo, Municipality of Triacastela. Individuals were captured with snap traps (Supercat® Swissinno, Switzerland) placed in galleries and activated during one day by local farmers. Specimens were transported to the Anatomy Department of the Faculty of Veterinary Medicine to conduct necropsies in no case taking more than two hours. Fossorial water voles are considered a key pest species in grasslands and their demographic densities need to be controlled in accordance with article 15 of the Law 43/2002 of plant health (BOE 2008). The recommendations of the Directive of the European Parliament and the Council on the Protection of Animals Used for Scientific Purposes (Directive 2010/63/UE 2010) were considered in all procedures.

### 2.2 Samples dissection

#### Nasal cavity

The study of the nasal cavity was conducted with the aim of macroscopically visualizing the VNO in situ and examining its topographical anatomy, giving additional attention to the functionality of the incisive duct. To extend this study to the microscopic level, complete decalcification of the nasal cavity was required in order to cut it with the microtome. For this purpose, the nasal cavity was transversely sectioned with a saw at the level of the molars and immersed in a decalcifying solution (Shandon TBD – 1 Decalcifier; Thermo, Pittsburgh, PA) for three days under agitation. Subsequently, it was washed in running water for 5 hours, and then fixed in Bouin’s fluid, replaced after 24 hours with 70% ethanol, and later embedded in paraffin. The block was serially sectioned into 6 µm thick sections, from the caudal end of the sample to the opening of the incisive duct into the oral cavity.

#### Vomeronasal organ and nerves

To avoid the negative effect of decalcification on immunohistochemical techniques, it is preferable to process the VNO after performing a meticulous dissection in which all remnants of bone tissue are removed. This allows its sectioning with the microtome without requiring decalcification. To do this, the nasal and maxillary bones were removed, providing access to the nasal cavity. After removing the dorsal and ventral nasal conchae, the VNO is observed protruding beneath the mucosa of the nasal cavity, lateral to the vomer bone. With extreme care the VNO was extracted, separating it from the vomer bone using forceps. Based on available information about the VNO of other rodents (Vaccarezza et al. 1981; Taniguchi and Mochizuki 1982; Oikawa et al. 1994), it was expected that the VNO of the water vole would have a capsule of bony nature. For this reason, an additional step was required; with the help of a surgical microscope (Zeiss OPMI 1), this bony capsule was removed, preserving the integrity of the delicate duct and vomeronasal parenchyma. This procedure was carried out on 4 animals. The extracted VNOs were fixed in Bouin’s liquid, embedded in paraffin, and serially sectioned in the transverse plane for histological study.

### 2.3 Processing of samples for microscopic study

#### Paraffin embedding

This was done gradually, immersing the samples progressively in distilled water, 70% ethanol, 90% ethanol, 96%° ethanol, and pure ethanol (3 baths). They were then cleared with an ethanol-xylene (1:1) mixture and two washes in xylene. After this bath, the tissue becomes permeable to the paraffin in which it is submerged at 60°C for a minimum of 3 hours. The sample was then covered with paraffin using a dispenser to form a solid block.

#### Sectioning

Sections were made using a rotary microtome with a thickness of 6–7 µm, serially sectioning all samples in the transverse plane along their entire length from caudal to cranial.

### 2.4 General histological stainings

Hematoxylin - eosin (HE) staining was used to conduct a general study of the structures. For a detailed study and to highlight certain tissue components Alcian Blue Staining (AB) was used This technique is used to mark among the acid mucopolysaccharides, which are stained with blue color.

### 2.5 Immunohistochemical stainings

The protocol is described in detail in Ortiz-Leal et al. (2023). As a first step, it is necessary to block endogenous peroxidase to ensure that the development is specific at the primary antibody binding site. To achieve this, the slide is incubated in a 3% hydrogen peroxide solution in distilled water, which depletes the reserves of endogenous peroxidase, deactivating it. After three washes in 0.1 M phosphate buffer (PB) pH 7.2, the blocking of non-specific bindings is required. For this, a blocking serum is added from the same species from which the secondary antibody is derived (ImmPRESS TM Anti-Rabbit and Anti-Mouse Kit from Vector Labs, Burlington, USA). The blocking was done at room temperature for a minimum of 20 minutes.

Primary antibodies employed (Table 1):

**Table 1.**
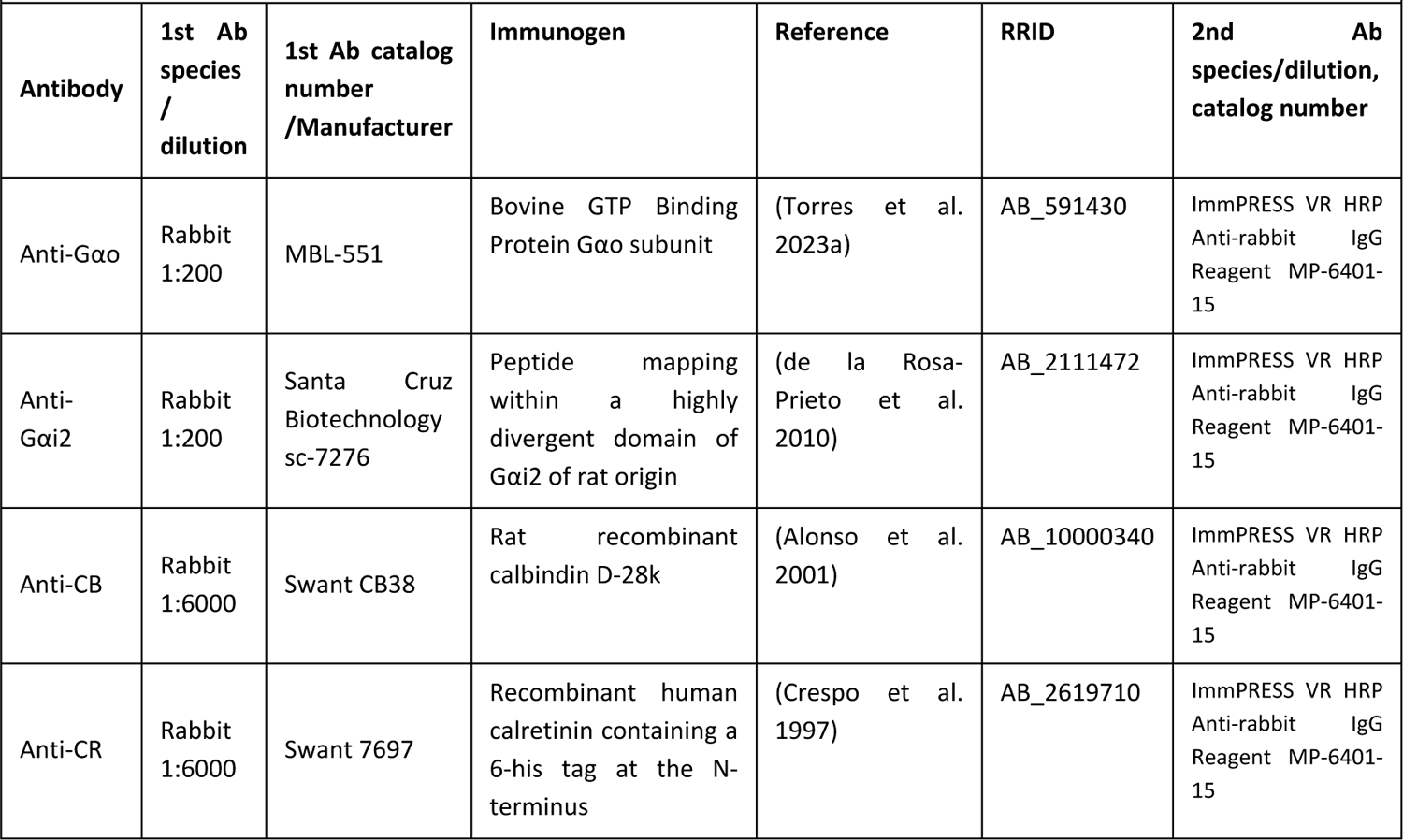
Detailed information on the antibodies used in this study: Species of elaboration, dilution, catalogue number, manufacturer, target immunogens, relevant reference for each antibody, RRID codes, and secondary antibody employed.

#### Anti - Gαi2

This antibody specifically binds to the i2 family of the α subunit of the G protein, belonging to the transduction cascade. It indicates the distribution of the expression of the vomeronasal genes from the V1R receptor family (Suárez et al. 2011; Pallé et al. 2020).

#### Anti - Gαo

This antibody specifically binds to the olfactory family or α subunit of the G protein. In this case, it indicates the distribution of the expression of the V2R receptor genes (Francia et al. 2015).

#### Anti - calbindin (CB)

Antibody against calbindin, a protein that is part of the superfamily of cytoplasmic calcium-binding proteins, which play a regulatory role in calcium (Baimbridge et al. 1992).

#### Anti - calretinin (CR)

Antibody with specific binding against calretinin, a protein that is part of a family of cytoplasmic calcium-binding proteins (Alonso et al. 2001).

Antibodies targeting calcium-binding proteins, are useful distinguishing neuronal subtypes in rodents (Abe et al. 1992; Kishimoto et al. 1993; Bastianelli and Pochet 1995; Fujiwara et al. 1997; Jia and Halpern 2003) and lagomorphs (Villamayor et al. 2020).

After overnight incubation at 4°C, samples were washed three times in PB, and then, depending on the blocking agent used, were incubated for 30 min with either the ImmPRESS VR Polymer HRP Anti-Rabbit IgG or Anti-mouse IgG reagents. Prior to the visualization stage, all samples were rinsed for 10 minutes in 0.2 M Tris-HCl buffer at pH 7.61. DAB chromogen was used for visualizing. A 0.003% hydrogen peroxide and 0.05% 3,3-diaminobenzidine (DAB) solution in a 0.2 M Tris-HCl buffer were used to visualize the ensuing reaction, which resulted in a brown-colored deposit. Negative controls omitted the primary antibodies.

### 2.6 Lectin histochemical labeling

Lectin labeling, although similar to immunohistochemical techniques, uses specific proteins called lectins. These lectins possess domains capable of recognizing and non-covalently binding to terminal sugars in tissues, forming glycoconjugates. However, lectins do not have an immune origin (Leathem and Atkins 1983). In this study, we worked with two lectins: UEA and LEA. The protocol is described in detail in Ortiz-Leal et al. (2022a).

#### *Ulex europaeus* (UEA)

It is an agglutinin derived from gorse (*Ulex europaeus*) that primarily recognizes terminal L-fucose belonging to glycoproteins and glycolipids. UEA was utilized due to its role as a specific vomeronasal marker in species including dogs (Salazar et al. 1994), foxes (Ortiz-Leal et al. 2020), and wolves (unpublished observations).

#### Lycopersicum esculentum (LEA)

LEA has a maximum affinity for N-acetylglucosamine. LEA has served as a comprehensive marker for both the MOS and VNS in rodents such as rats (Salazar and Sánchez Quinteiro 1998), mice (Keller et al. 2022) capybaras (Torres et al. 2020), and carnivores like minks (Salazar et al. 1998) and meerkats (Torres et al. 2021).

The protocol for LEA lectin is slightly different from the protocol followed for UEA. Both lectins undergo prior blocking of non-specific bindings with 2% bovine serum albumin (BSA). After this incubation, in the case of the UEA labeling protocol, the sample is incubated with the pure lectin (Vector L – 1060) at a 1:60 ratio for 1 hour, after which a commercially peroxidase-linked antibody against UEA is added (DAKO, P289). For LEA labeling, a biotinylated lectin (Vector B – 1175) at 20 µg/ml is used, which is incubated overnight. After this step, it is necessary to incubate the sample for 90 min in an ABC complex, which consists of an avidin-biotin complex and peroxidase. During the incubation, this complex binds to the lectin, allowing the peroxidase reaction. In both cases, the development is identical to that explained in the immunohistochemistry protocol. As controls, tests without lectin addition and also pre-absorbed lectins with excessive corresponding sugars, were performed.

### 2.7 Image acquisition

Digital photographs were taken using a digital camera from Karl Zeiss, model MRc5 Axiocam, connected to a microscope of the same brand, model Axiophot. Given the large size of the photographed areas and to achieve maximum definition, the presented photos are the result of combining a mosaic of photos (ranging in number from 6 to 90) using an automatic fusion program (PTGui, Rotterdam, Netherlands). The Adobe Photoshop CS4 software (Adobe Systems, San Jose, CA) was used for those images that required digital enhancement of brightness, contrast, and white balance. However, no enhancements, additions, or relocations of the image features were made.

## RESULTS

Due to the limited dimensions of the fossorial water vole nasal cavity, macroscopic external visualization of its VNO is challenging. Consequently, we have described its key anatomical features and most significant topographical relationships based on the decalcified histological series. Nonetheless, a detailed observation of the ventral part of the palate reveals an elevation of the mucosa corresponding to the incisive papilla (Fig. 1). The significance of this papilla lies in its role as the opening point for the nasopalatine duct, which connects the oral cavity to the nasal cavity and simultaneously allows chemical signals to access the vicinity of the VNO. The presence of a wide philtrum on the upper lip (Fig. 1A) is crucial for facilitating the passage of molecules into the incisive area. The incisive papilla is characterized by its triangular shape (Fig. 1B) and occupies the entire anterior end of the palate (Fig. 1C).

**Figure 1.**
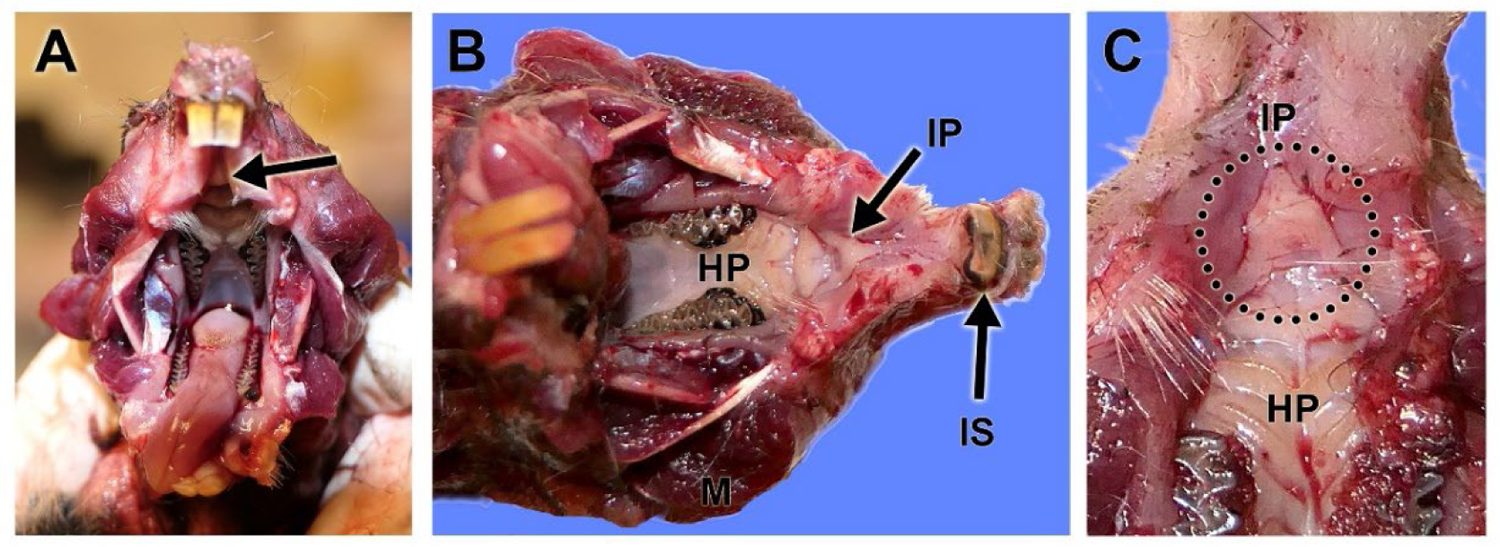
Macroscopic anatomy of the water vole incisive papilla. **A.** Rostroventral view of the palate showing the pathway (arrow) from the philtrum towards the incisive papilla (IP). **B.** Ventral view of the hard palate (HP) showing the triangular shape of the IP. **C.** Closer look to the IP area (dotted circle). M: Maseter muscle; IS. Incisor teeth.

For the general microscopic description of the VNO, we will refer to a decalcified transverse section of the nasal cavity taken from the central part of the VNO rostrocaudal axis (Fig. 2). At this level, the VNO is located in the ventral part of the cavity, occupying almost the entire ventral meatus. Both organs rest on the vomer bone, which emits two lateral projections and central one that completely encapsulates the soft tissue of the organ. Thus, the VNO is composed of a solid bone capsule and a parenchyma or soft tissue, consisting of glands, blood vessels (predominantly veins), and nerves, which enclose the vomeronasal duct (VD) (Fig. 2B). The VD is crescent-shaped, with its medial part lined by a pseudostratified columnar sensory neuroepithelium and its lateral part by a pseudostratified respiratory epithelium.

**Figure 2.**
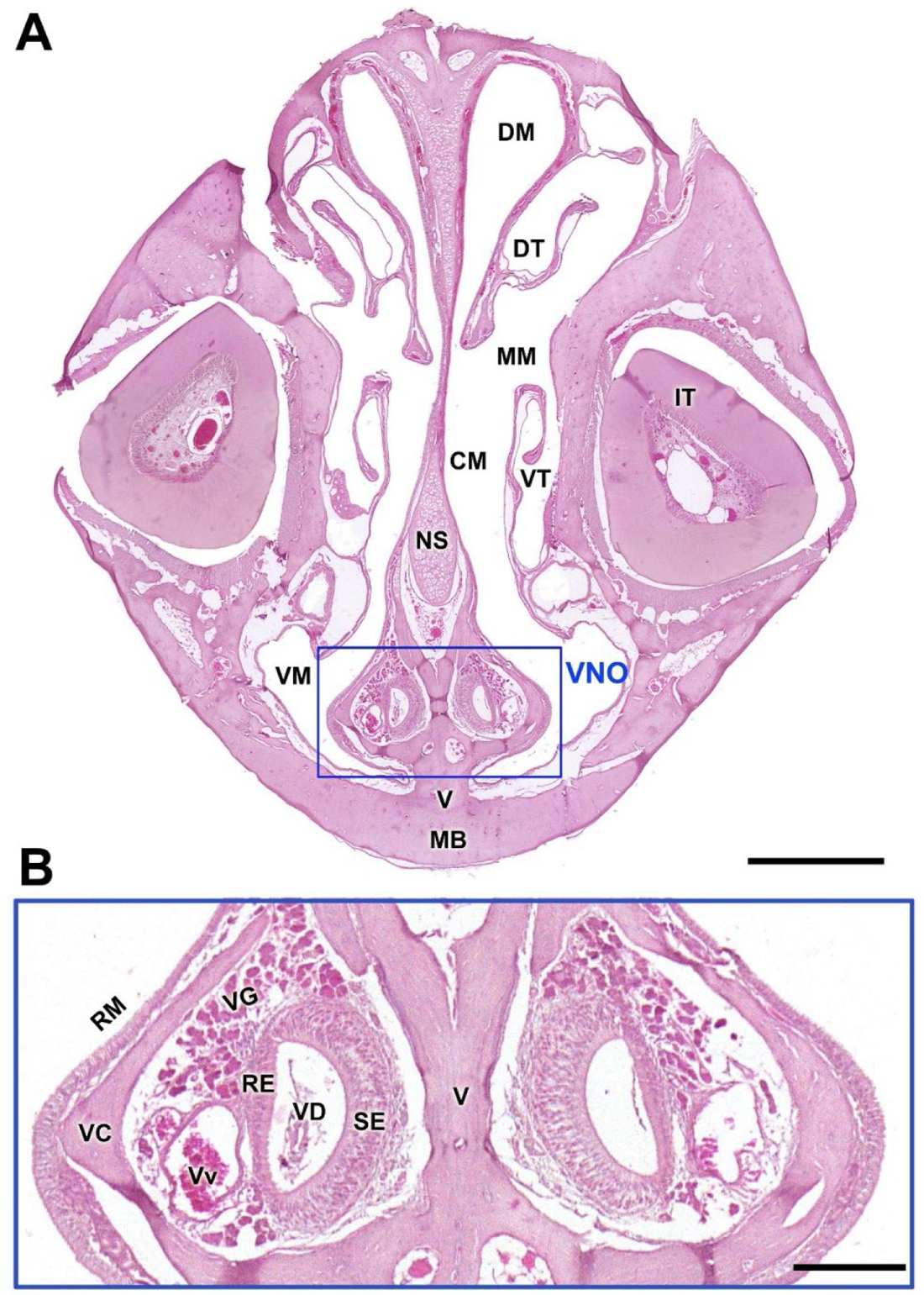
Histological study of the nasal cavity of the water vole showing the topographical anatomy of the VNO. **A.** Decalcified cross-section of the nasal cavity at the level of the central part of the VNO. The vomeronasal organs are located on both sides of the base of the nasal septum (NS) encapsulated by the two lateral plates of the vomer bone (V), occupying almost entirely the ventral meatus (VM) of the nasal cavity. **B.** Magnification of the box shown in A where the main components of the VNO can be identified. Hematoxylin-eosin staining. DM: Dorsal meatus, DT: Dorsal turbinate, IT: Root of the incisor tooth, MB: Maxillary bone, MM: Medium meatus, RE: Respiratory epithelium, RM: Respiratory mucosa, SE: Sensory epithelium, VC: Vomeronasal capsule, VD: Vomeronasal duct, VG: Vomeronasal glands, VT: Ventral turbinate, Vv: Veins. Scale bars: (A) 1mm, (B) 250 µm.

Serial histological sections of the VNO allowed us to confirm significant differences in its configuration along its rostrocaudal axis. These are depicted in Fig. 3. A particularly important feature is the connection of the VD to the exterior, through two small meatuses located at the anterior end of the nasal cavity (Fig. 3B). At this point, the VD is lined by a simple columnar epithelium, with a reduced development of the bone capsule. However, it is complemented by the presence of a paraseptal cartilage. Progressing caudally, there is a marked increase in the development of the bone capsule, especially medially, as the paraseptal cartilage gradually disappears (Fig. 3D). At this level the medial epithelium of the VD is sensory in nature. In the VNO central portion (Fig. 3A, E), all soft tissue components reach their maximum expression. The development of glandular tissue from this point on is remarkable, with tubuloacinar glands spanning the entire lateral part of the parenchyma and opening at the dorsal commissure of the VD (Fig. 3E).

**Figure 3.**
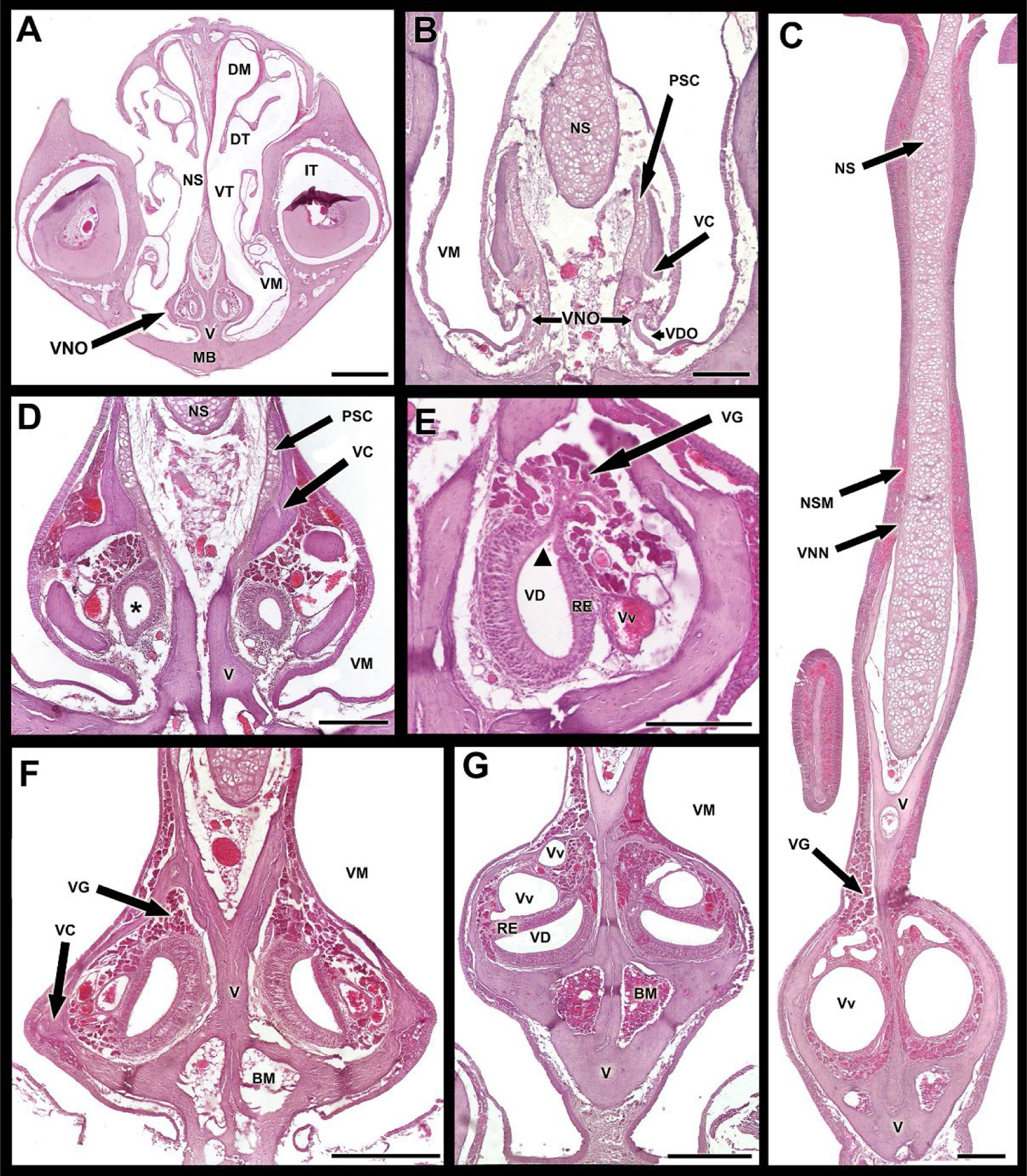
Serial histological study of the water vole VNO displaying the structural changes along its rostro-caudal axis. **A.** Cross-section at the central level of the VNO. **B.** Cross-section of the base of the nasal septum (NS) in the rostral part of the nasal cavity (NC) showing the opening of the vomeronasal duct (VDO) in the ventral meatus (VM). **C.** Complete cross-section of the NS in the caudal part of the nasal cavity displaying the two large nasopalatine veins (Vv) supplying the VNO accompanied by glandular tissue (VG). **D.** Cross-section of the base of the NS in the anterior part of the NC demonstrating how the bony capsule (VC) of the vomer bone (V) incompletely encloses the vomeronasal duct (asterisk). **E.** Transverse section of the VNO showing the opening of the vomeronasal glands (VG) in the dorsal commissure of the duct (VD). **F and G.** Cross-sections at the caudal levels of the vomeronasal organ showing how the vomeronasal duct rotates in a lateroventral direction. Hematoxylin-Eosin staining. BM: Bone marrow; DM: Dorsal meatus, DT: Dorsal turbinate, IT: Incisor tooth, MB: Maxillary bone, NSM: Nasal septum mucosa; RE: Respiratory epithelium, PSC: Paraseptal cartilage, VNN: Vomeronasal nerves; VT: Ventral turbinate. Scale bars: (A) 1000 µm, (B) 50 µm, (C,D) 250 µm, (E) 100 µm, (F, G) 500 µm.

Caudally, the vascular component occupies a substantial part of the parenchyma, displacing the glandular component (Fig. 3F-H). It consists mainly of a large vein that is most pronounced in the most caudal level depicted (Fig. 3H). At this level, the VD is no longer present as it ends blindly, but the glandular tissue and the thick bony capsule enclosing the organ still persist. It is notable that the VD, while maintaining its crescent shape to its most caudal end, undergoes a significant rotation such that its medial sensory component becomes ventral, and the lateral respiratory component becomes dorsal (Fig. 3G).

The vomeronasal nerves (Fig. 3H) are formed by the confluence of axons from the neuroepithelium. In the fossorial water vole, they are of small caliber, and their visualization is enhanced by immunohistochemical techniques. Alcian blue staining was used to determine the nature of the glandular tissue secretion (Fig. 4). The mainly negative result (Fig. 4A, C) suggests the neutral nature of the mucopolysaccharides secreted by the vomeronasal glands. Only at the most caudal level is a small number of alcian-blue positive glands observed (Fig. 4B).

**Figure 4.**
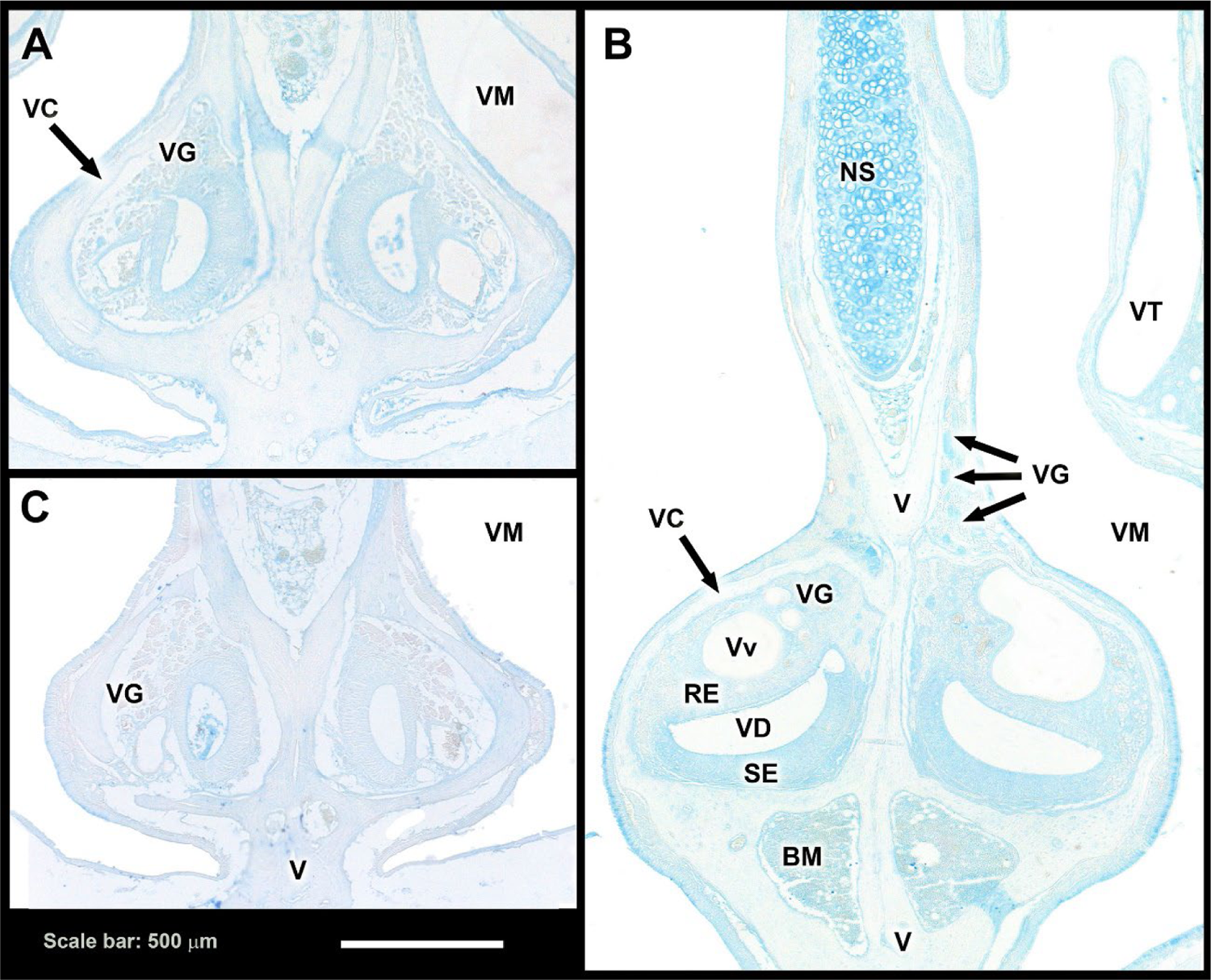
Fossorial water vole VNO. Study on the secretion of acidic mucopolysaccharides. Alcian blue staining. **A and C.** Decalcified cross-sections of the VNO in its central (A) and rostral (C) levels show the absence of acidic polysaccharide secretion. **B.** In the caudal levels of the organ, a sparse number of AB-positive vomeronasal glands (VG) are observed. BM: Bone marrow, NS: Nasal septum, RE: Respiratory epithelium, SE: Sensory epithelium, V: Vomer bone, VC: Bony vomeronasal capsule, VD: Vomeronasal duct, VM: Ventral meatus, VT: Ventral concha, Vv: Veins.

### Lectin histochemical study

UEA lectin (Fig. 5) identified the following components of the vomeronasal organ: the two epithelia that form the vomeronasal duct, the vomeronasal glands, and the vomeronasal nerves (Fig. 5A). The labeling in the sensory epithelium comprised the dendrites and somata of the neuroreceptor cells, and the vomeronasal axons in the lamina propria. In the respiratory epithelium, UEA labeled the superficial mucomicrovillar complex and the nuclei of apical cells.

**Figure 5.**
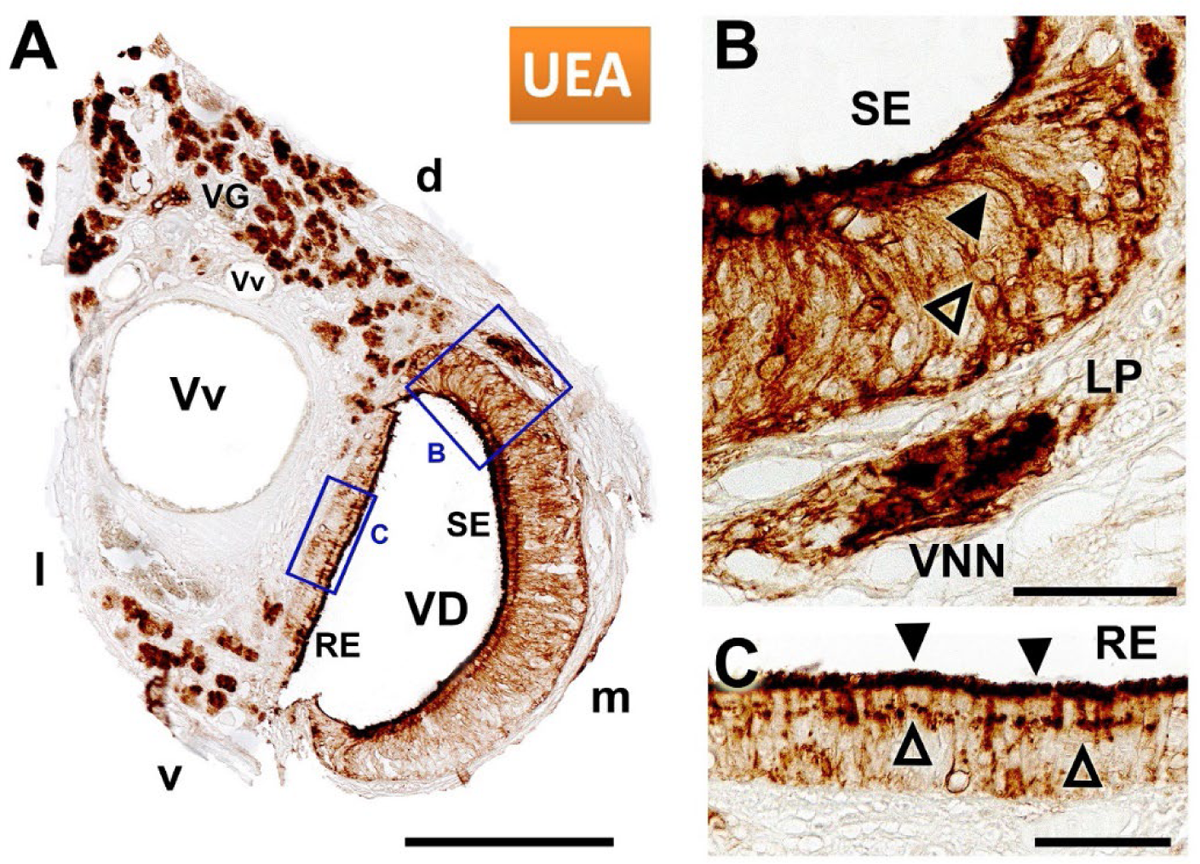
UEA lectin histochemical study of the fossorial water vole VNO. **A.** Vomeronasal duct (VD) labeling showing positivity in the glandular tissue (VG), the vomeronasal sensory epithelium (SE) (enlarged in B), and the respiratory epithelium (enlarged in C). **B.** The labeling in the SE comprised dendrites (black arrowhead), neuronal somata (hollow arrowhead), and vomeronasal nerves (VNN) in the lamina propria (LP). **C.** In the respiratory epithelium (RE), UEA labels the surface of the mucomicrovillar complex (black arrowhead) and the nuclei of apical cells (hollow arrowhead). d: Dorsal, l: Lateral, m: Medial, v: Ventral; Vv: Veins. Scale bars: (A) 250 µm (B & C) 50 µm

The LEA lectin (Fig. 6) shows a different pattern. In particular, staining in the neuroepithelium is less intense (Fig. 6A). It is primarily localized in the somas of neuroepithelial and basal cells and to the mucomicrovillar complex, whereas dendritic processes stain weakly (Fig. 6B). Labeling in the respiratory epithelium is confined to the mucous complex which also overlaps this epithelium (Fig. 6C).

**Figure 6.**
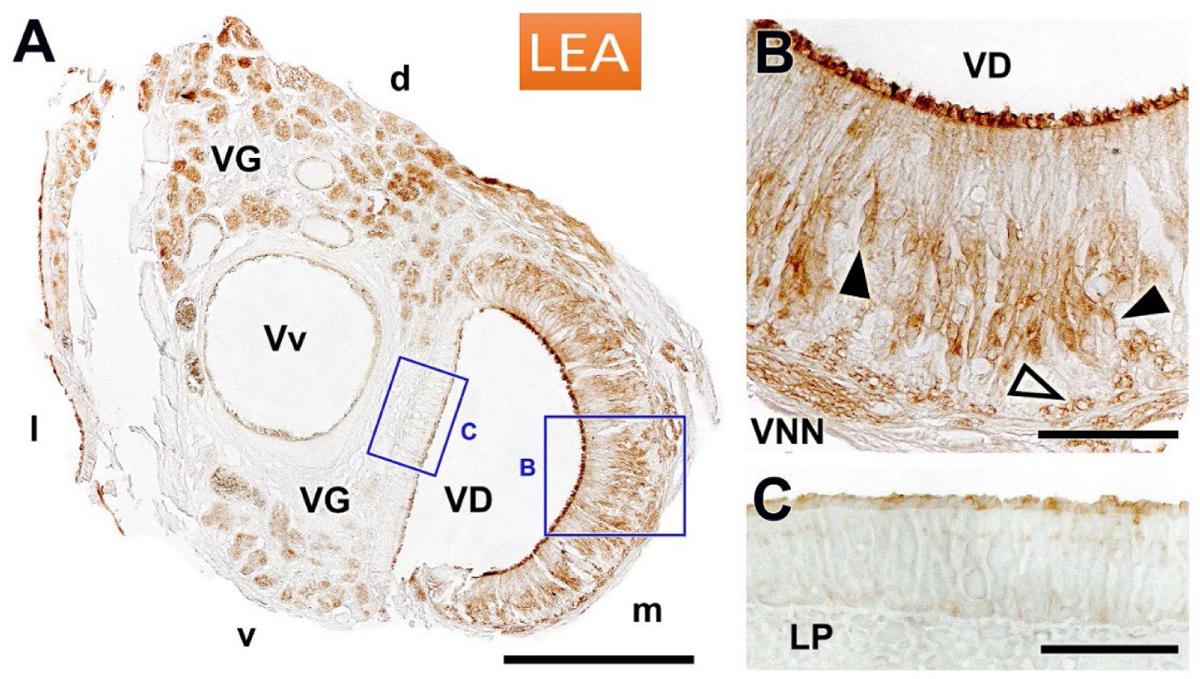
LEA lectin histochemical staining of the water vole VNO. **A.** Labeling of the vomeronasal duct (VD) showing positivity in the glands (VG), in the vomeronasal sensory epithelium (enlarged in B), and in the respiratory epithelium (enlarged in C). **B.** The labeling in the sensory epithelium was localized at the luminal border, in the neuronal somata (black arrowhead), and in the basal cells of the epithelium (hollow arrowhead). **C.** In the respiratory epithelium, LEA labels very weakly the surface of the mucomicrovillar complex. d: Dorsal, L: lateral, LP: Lamina propria, m: Medial, v: Ventral; Vv: Veins. Scale bars: (A) 250 µm, (B,C) 50 µm.

### Immunohistochemical study

The immunohistochemical study focused on the use of antibodies against the two subunits of the G proteins, Gα0 and Gαi2 (Fig. 7, 8), and two calcium-binding molecules, calbindin and calretinin (Fig. 9, 10). All of these are widely used in the morphofunctional characterization of the VNO.

**Figure 7.**
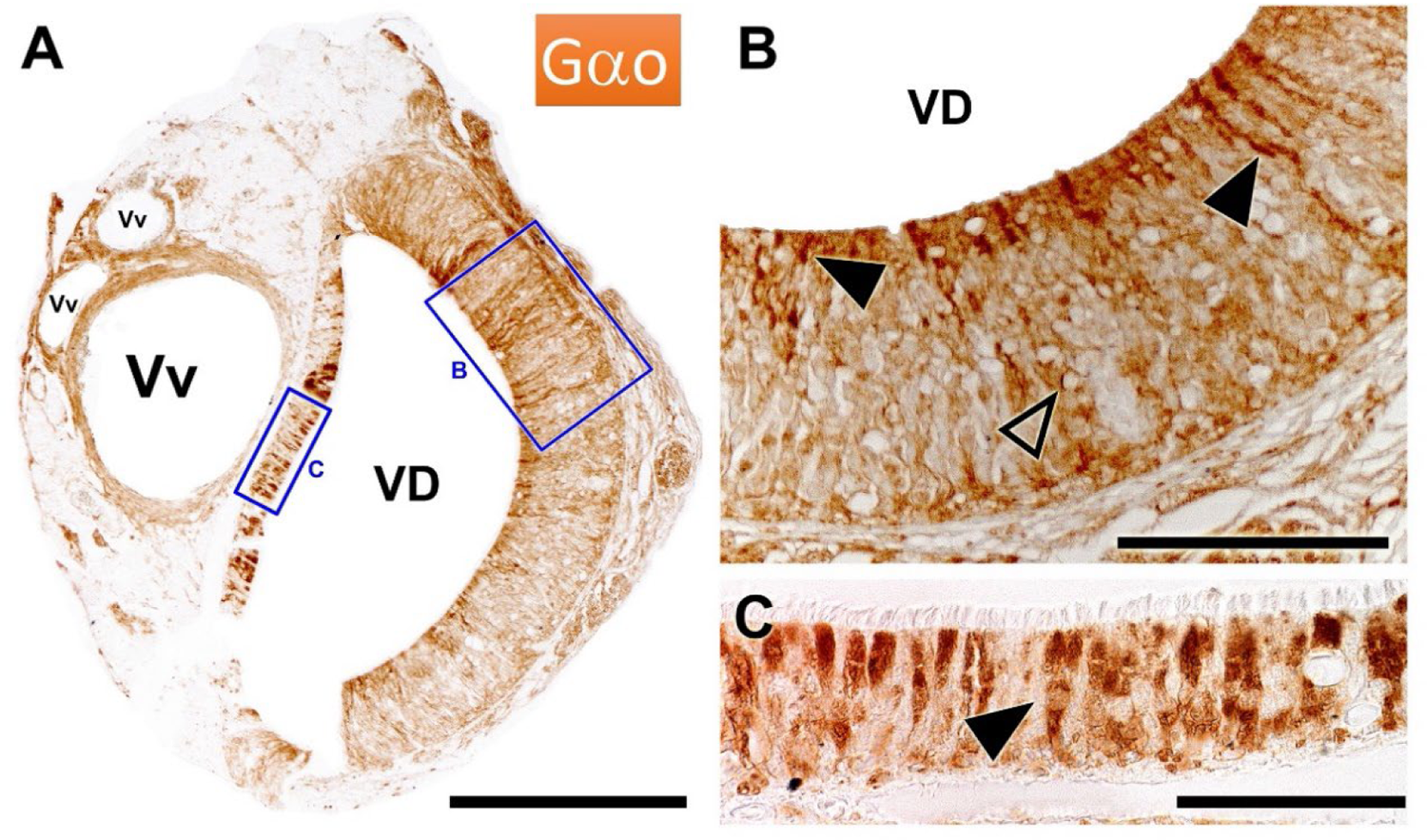
Immunohistochemical labelling of the fossorial water vole VNO with anti-G*α*o. **A.** Immunolabeling of the vomeronasal duct (VD) showing immunopositivity in the sensory epithelium (enlarged in B) and in the respiratory epithelium (enlarged in C). **B.** The labeling in the sensory epithelium is localized at the luminal border, in the dendritic knobs (black arrowhead), and in the neuroreceptors cells of the epithelium (hollow arrowhead). **C.** In the respiratory epithelium, epithelial cells with an enlarged immunopositive apical end are predominantly labeled (arrowhead). VD: Vomeronasal duct.; Vv: Veins. Scale bars: (A) 250 µm, (B) 100 µm, (C) 50 µm.

The antibody against Gαo produced intense labeling within the neuroepithelium of the VNO (Fig. 7A). At higher magnification, labeling was observed to be mainly concentrated in the neuronal dendrites and somata (Fig. 7B). Notably, in the respiratory epithelium, individual cells were stained intensely (Fig. 7C). The positivity to this marker suggests their chemosensory nature.

The antibody against Gαi2 stained exclusively the sensory epithelium (Fig. 8A), albeit in a restricted pattern, predominantly labeling the apical neurons (Fig. 8B).

**Figure 8.**
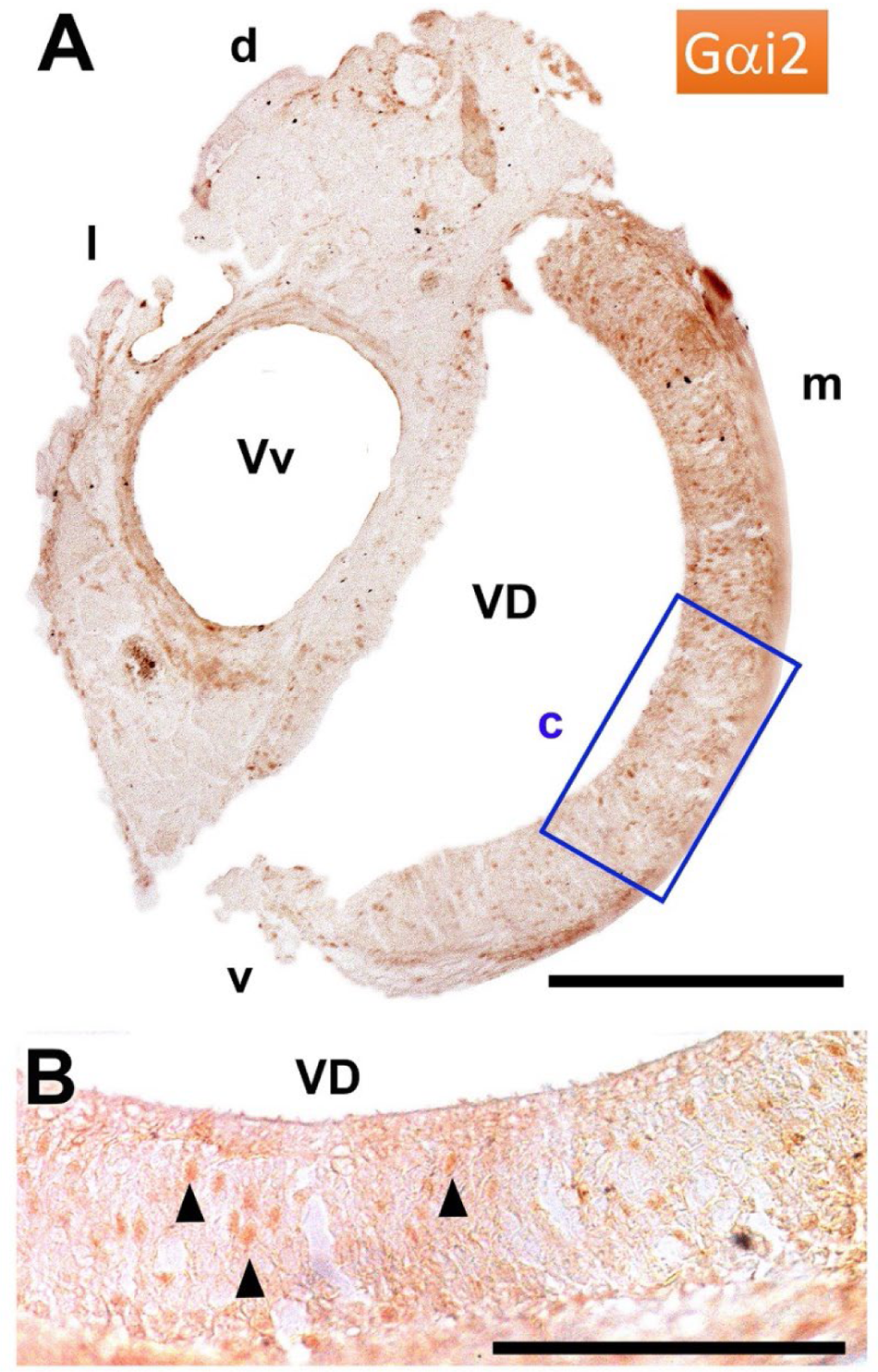
Immunohistochemical labelling of the fossorial water vole VNO with anti-G*α*i2. **A.** Immunolabelling showing immunopositivity to Gαi2 in the sensory epithelium of the vomeronasal duct (VD) (enlarged in B). **B.** Immunopositive neuroreceptor cells of the sensory epithelium (arrowhead) show a random distribution, although predominantly apical. d: Dorsal, l: Lateral, m: Medial, v: Ventral; Vv: Veins. Scale bars: (A) 250 µm, (B) 500 µm, (C) 100 µm.

Calbindin is exclusively present in all neural elements of the vomeronasal pathway, producing intense labeling within the neuroepithelium of the organ (Fig. 9 A). The immunolabeling in the sensory epithelium encompasses the dendrites, somata, and the vomeronasal nerves in the lamina propria. The intensity of the labeling clearly delineates the presence of intraepithelial capillaries.

**Figure 9.**
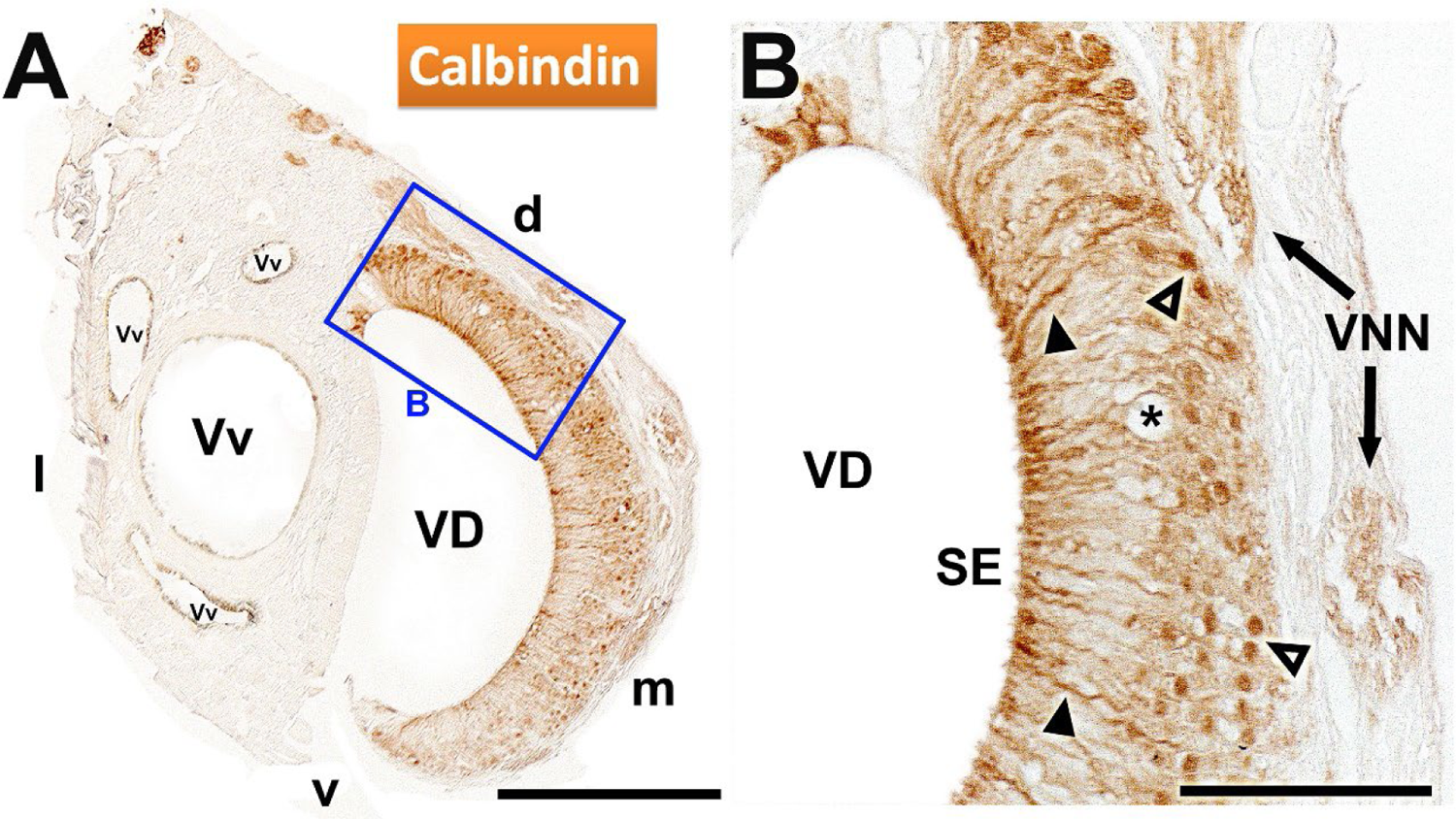
Immunohistochemical labeling of the water vole VNO with anti-calbindin. **A.** Immunostaining of the vomeronasal duct (VD) produces immunopositivity in the sensory epithelium (enlarged in B) and in the vomeronasal nerves. **B.** The anti-CB immunolabeling in the sensory epithelium (SE) encompasses the dendrites (black arrowhead), somata (hollow arrowhead), and the vomeronasal nerves (VNN) in the lamina propria. It is remarkable the presence of intraepithelial capillaries (asterisk). d: Dorsal; l: Lateral; m: Medial; v: Ventral; Vv: veins. Scale bars: (A) 250 µm, (B) 100 µm.

Calretinin shows a similar pattern, although with greater intensity (Fig. 10A,B). Since the immunohistochemical labeling was performed on consecutive sections, a higher density of immunopositive somas can apparently be observed with this marker compared to calbindin.

**Figure 10.**
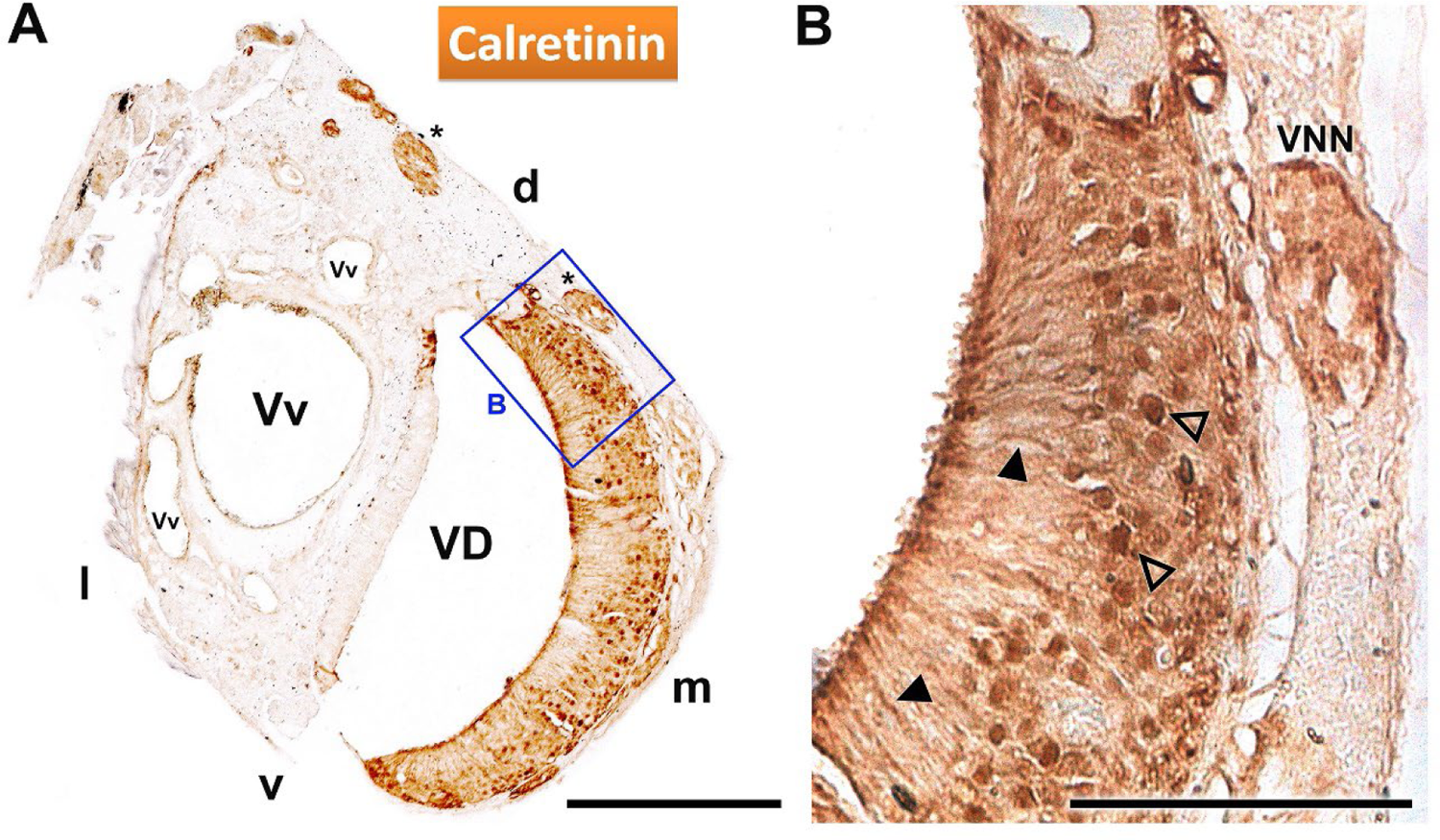
Immunohistochemical labeling of the water vole VNO with anti-calretinin. **A.** Immunostaining of the vomeronasal duct (VD) showing positivity in the sensory epithelium (enlarged in B) and in the vomeronasal nerves (asterisk). **B.** The immunolabeling in the sensory epithelium encompasses the dendrites (black arrowhead), somata (hollow arrowhead), and the vomeronasal nerves in the lamina propria (VNN). d: Dorsal, l: Lateral, m: Medial, v: Ventral; Vv: veins. Scale bars: (A) 250 µm, (B) 100 µm.

## DISCUSSION

While the use of signals involved in pheromone-mediated chemical communication has been shown to be instrumental in the invertebrate pests control (Reddy and Guerrero 2010; Rizvi et al. 2021), the application of pheromones in mammalian pest control remains relatively nascent (Shapira et al. 2013; Clapperton et al. 2017). Part of the challenge stems from a lack of anatomical information, which would clearly delineate the morphological and functional basis of the main system responsible for detecting chemical cues—the vomeronasal system. Although there is a substantial amount of information available on the vomeronasal system of mammals (Wysocki 1979; Halpern 1987; Halpern and Martínez-Marcos 2003; Torres et al. 2023b), we are currently experiencing a situation in which genomic knowledge has outpaced its morphological counterpart. There is a discernible disparity between these two domains, with geneticists themselves calling for anatomical information on which to base their genomic or transcriptomic observations (Zhang and Nikaido 2020; Yohe and Krell 2023). Bridging this gap requires a renewed focus on morphological foundations.

To further complicate the matter, the vomeronasal system, given its extensive morphological diversity among mammalian groups, makes extrapolation from one species to another problematic. Such extrapolations are not only risky across different mammalian orders but also within the same family. For example, comparing laboratory rodents, such as rats and mice, with wild rodent species reveals distinct and species-specific characteristics. Rodents such as capybara (Torres et al. 2020) and chinchilla (Oikawa et al. 1994) show vomeronasal systems distinct from rats (Vaccarezza et al. 1981; Salazar and Sanchez-Quinteiro 1998) and mice (Barrios et al. 2014a) in terms of sensory epithelium organization, neurochemical patterns, maturation rate, neuroregenerative capabilities, among other features. These may play a key role in understanding the impact of pheromones and kairomones on behavior and reproductive physiology.

Chemical signaling can be exploited as a potential tool within sustainable control methods aimed at containing fossorial water vole populations. This is underscored by recent comprehensive studies using proteomic techniques and chromatography to identify chemicals characterizing some of their exocrine secretions (urinary, lateral gland) (Nagnan-Le Meillour et al. 2019). Further, the attractive effects of some of these compounds have been analyzed both in lab settings and natural conditions (Poissenot et al. 2023). Such studies emphasize the importance of understanding the differentiation degree and functionality of the vomeronasal system in these species.

For this reason, we have aimed to thoroughly characterize the macroscopic, histological, and neurochemical features of the vomeronasal organ of the fossorial water vole to determine its differentiation and functionality levels. The following sections will discuss these aspects within the broader context of rodents.

The vomeronasal organ of the fossorial water vole exhibits a complex anatomy comparable to that of species with a higher degree of VNO differentiation. Externally, the small size of its nasal cavity makes it difficult to visualize the organ. However, we were able to assess the remarkable development of the mucosal elevation corresponding to the incisive papilla. This triangular-shaped structure occupies a significant portion of the anterior part of the palate roof, with a more pronounced development compared to other rodents, including the rat (Mahdy and Mohammed 2021), mouse (Navarro et al. 2017), capybara (Torres et al. 2020), and even lagomorphs (Villamayor et al. 2020). The papilla is pivotal as the opening point for the nasopalatine duct, which links the oral and nasal cavities and plays a crucial role in allowing chemical signals to approach the VNO. The presence of a wide philtrum on the upper lip enhances the papilla importance, facilitating the movement of molecules to the incisive area (Døving and Trotier 1998).

From a comparative point of view, the anatomical location of the rodent VNO in relation to the nasal septum shows variations. For instance, the VNO is distinctly situated at the base of the nasal septum, dorsally to the vomer bone in rodents like the rat (Weiler 2005) and mouse (Ibarra-Soria et al. 2014). On the contrary, it is located on the prominent palatine process of the incisive bone in others such as capybara (Torres et al. 2020) and chinchilla (Jurcisek et al. 2003). Histological series of the water fossorial vole nasal cavity suggest this species follows the same model as the rat and mouse. The fossorial water vole also shares with both species a similar configuration of the VNO capsule, which consists of a bone capsule with two lateral projections and a central one, completely surrounding the VNO soft tissue (Vaccarezza et al. 1981; Salazar et al. 2016). This encapsulation serves as a protective and supportive role for the vomeronasal pump function (Meredith and O’Connell 1979; Meredith et al. 1980; Salazar et al. 1995; Salazar et al. 2003). This model is not the only one observed in rodents as various families exhibit a range of structural possibilities. The capybara has a rostrally cartilaginous capsule that transitions to bone caudally (Torres et al. 2020), whereas the naked mole rat, *Heterocephalus glaber*, features an exclusively cartilaginous capsule (Dennis et al. 2020). Meanwhile, two fossorial mole rat species from the Bathyergidae family, *Cryptomys hottentotus* and *Fukomys damarensis*, show a medially cartilaginous and laterally bony envelope (Dennis et al. 2020).

The histological study of the fossorial water vole VNO demonstrates the intricate configuration of the rostral part of the organ and confirms the connection of the vomeronasal duct to the outside via a small meatus, similar to what happens in the rat (Vaccarezza et al. 1981; Salazar and Sanchez-Quinteiro 1998) and mouse (Breipohl et al. 1981). While there is limited glandular tissue development at this level, in the VNO central sections, there is a pronounced presence of glandular tissue. This differs from what was observed in rat and mouse, which have at that level a reduced presence of glandular tissue (Mendoza 1986; Barrios et al. 2014a). For the fossorial water vole, the glandular tissue, located dorsally in the parenchyma of the rostral levels, progressively extends ventrally and laterally in caudal sections. Proper glandular secretions are vital for maintaining an optimal environment within the VNO, such as pH and moisture level (Takami et al. 1995). Moreover, these secretions dissolve and transport pheromones to the VNO sensory receptors and might modulate pheromone activity (Zufall and Munger 2001). The remarkable development of the glandular tissue in the fossorial water vole suggests an adaptation to its subterranean lifestyle, which would need an especially abundant glandular secretion.

We expanded our investigation of the glandular system by applying alcian blue staining and histochemical labeling with lectins. Our study, which showed only a few alcian-blue positive glands at the most caudal level, confirms that the glandular secretion is mostly neutral. This finding is consistent with most studied rodents such as the rat (Salazar and Sanchez-Quinteiro 1998), guinea pig (Pastor et al. 1992), mouse (Cuschieri and Bannister 1975; Kondoh et al. 2020), vole (Roslinski et al. 2000), hamster (Taniguchi and Mochizuki 1982), and chinchilla (Oikawa et al. 1994). Only the capybara has a high proportion of alcian blue-positive glands (Torres et al. 2020).

In the fossorial water vole, other aspects, such as the vascular system organization and the vomeronasal duct epithelial lining, show an organization similar to that in the rat and mouse. In terms of vasculature, it features a large central vein responsible for venous return. For the fossorial water vole, it is striking how, towards the VNO caudal end, a large part of the parenchyma is almost entirely occupied by this vein. In terms of epithelial organization, it aligns with the patterns observed in both the rat and mouse, characterized by a semilunar shape and differentiation between a thick medial sensory epithelium and a thin lateral respiratory epithelium. Interestingly, in the fossorial water vole, the vomeronasal duct undergoes a prominent caudal rotation. It would be intriguing to determine if this has any functional significance. Finally, the pronounced presence of intraepithelial capillaries suggests a promising pathway for future studies.

### Lectin histochemical and immunohistochemical stainings

Our results highlight distinct patterns that can provide deeper insights into the morphofunctional architecture of this organ. The two lectins, UEA and LEA, stained the vomeronasal glands, with UEA showing stronger intensity both dorsally and ventrally in the parenchyma. Both lectins have distinct specificities: L-fucose for UEA (Kondoh et al. 2017) and N-acetyl-glucosamine for LEA (Plendl and Sinowatz 1998). This emphasizes the broad diversity of glycoconjugates expressed in the VNO of the fossorial water vole. This contrasts with findings observed in rats (Salazar and Sánchez Quinteiro 1998) and mice (Keller et al. 2022), where the vomeronasal glands are UEA positive but LEA negative.

In the sensory vomeronasal epithelium of the fossorial water vole, both lectins, especially UEA, label the dendrites, somata, and axons of the neuroreceptor cells. This suggests that the glycoconjugates detected by them play a key role in recognizing and processing specific chemical signals. Both UEA and LEA also stain the sensory epithelium in rats (Lundh et al. 1989; Salazar and Sánchez Quinteiro 1998) and mice (Salazar and Sánchez Quinteiro 2003; Keller et al. 2022). However, a key difference exists compared to the fossorial water vole: there is a clear apical-basal zonation in the sensory epithelium in both species, especially in mice. In the fossorial water vole, neuroreceptor cells positive for both lectins are seemingly uniformly distributed throughout the sensory epithelium. In neither case does staining encompass all neuroreceptor cells, suggesting the possibility of complementary populations. To our knowledge, the only other rodent species studied with lectins is the capybara, which, like the fossorial water vole, does not exhibit this basal zonation (Torres et al. 2020). The significance of these differences remains unresolved.

Regarding the respiratory epithelium, it Is noteworthy how in the fossorial water vole the UEA lectin produces strong labeling, consistent with findings in mice with the same lectin (Lundh et al. 1989; Mendoza and Khünel 1991). Moreover, the lack of reactivity in the respiratory epithelium of the water vole against LEA aligns with observations by Keller et al. (2022) in the respiratory epithelium of mice.

The immunohistochemical analysis using antibodies specific to the G protein subunits, Gαo and Gαi2, holds particular interest. This is because the expression of both proteins in the VNO relates to the expression of the two main vomeronasal receptor families, V1R and V2R (Dulac and Axel 1995; Ryba and Tirindelli 1997). Immunopositivity to both markers in the sensory epithelium of the vomeronasal organ from the fossorial water vole confirms the functional expression of the two VR receptor families in this species. Anti-Gαo results in intense labeling, in dendrites and neuronal somata of the neuroreceptor cells, and within the vomeronasal nerves. Anti-Gαi2 displays a more restricted pattern within the global population of neuroreceptor cells, pointing to its role in a more specialized sensory signaling pathway. Studies in mice link V1R receptors with the detection of volatile pheromones and have associated them with aggressive and social behaviors in rodents (Chamero et al. 2011; Pallé et al. 2020). V2R receptors are involved in detecting non-volatile pheromones, often found in bodily fluids like urine, and are strongly associated with reproductive behaviors, such as mate recognition and estrous cycle synchronization among cohabiting females (Del Punta et al. 2002; Oboti et al. 2014).

It is significant that in the case of Gαo, we also found immunopositivity in individual cells within the respiratory epithelium, which highlights the potential chemosensory function of these cells. The morphology of these cells and their neurochemical pattern are comparable to the solitary chemosensory cells observed in mice (Ogura et al. 2010) and foxes (Ortiz-Leal et al. 2020).

A feature of particular interest concerning the expression of Gao and Gai2 proteins in the sensory neuroepithelium of the vomeronasal organ is the topographic organization of both cell populations. Specifically, it is known that in mice there is an apical-basal zonation such that neurons expressing Gαo are located in the basal part of the epithelium, while those expressing Gαi2 are situated in the apical region (Jia and Halpern 1996; Barrios et al. 2014a). Within rodents, this pattern has also been described for the rat (Jia and Halpern 1996; Takigami et al. 2000), although there are conflicting results for this species. As pointed out in the discussion of the VNO histochemical labeling with lectins, this apical-basal pattern in mice corresponds to an equivalent zonation discriminated by the UEA and LEA lectins (Salazar and Sánchez Quinteiro 2003). However, not all rodent species exhibit this zonation. This is the case for the capybara, a species in which both populations are interspersed within the neuroepithelium without following any discernible pattern (Torres et al. 2020). Our study in the fossorial water vole shows that this species follows this model. Not only do the two G proteins not follow a topographic pattern, but the LEA and UEA lectins also produce a diffuse labeling throughout the vomeronasal neuroepithelium. Beyond rodents, a diffuse pattern is found in species such as the wallaby (Torres et al. 2022) and rabbit (Villamayor et al. 2018). However, the apical-basal zonation has been described in the opossum (Halpern et al. 1995). All of this suggests that the topographic organization of the expression of the two VR receptor families follows a highly specific pattern whose significance remains to be elucidated.

The use of immunohistochemistry for calcium-binding proteins, namely calbindin and calretinin, further clarified the neural configuration within the fossorial water vole VNO. Both proteins are prevalent throughout all neural components of the VNO, underscoring their role in processing sensory signals. Differences in topography, both in terms of the number of cells expressing these proteins and their staining intensity, could provide further insights into the spatial aspects of odor detection. The higher intensity of calretinin immunostaining compared to calbindin in consecutive sections suggests a potential hierarchical or differential expression of these calcium-binding proteins. In both cases, the entire immunolabeled neuroreceptor cells are marked, highlighting their somas, dendritic processes, dendritic knobs, and axonal projections to the vomeronasal nerves. A similar pattern of intensity and density of marked neurons has been described in mice (Kishimoto et al. 1993), rats (Johnson et al. 1992; Jia and Halpern 2003), and capybaras (Torres et al. 2020).

In conclusion, the structural and neurochemical study of the fossorial water vole vomeronasal organ reveals a distinctly differentiated structure. This structure is equipped with all the necessary elements for performing chemosensory functions. The detailed staining patterns observed in both histochemical and immunohistochemical analyses further highlight the complex cellular and molecular configuration of the fossorial water vole VNO. The variation in staining with distinct markers suggests this organ may have multifunctional roles in sensory perception. Further research is necessary to accurately determine the specific roles of the characterized molecules, such as the lectin-detected glycoconjugates and calcium-binding proteins, which will enhance the understanding of the VNO function in sensory perception and its implications in the fossorial water vole behavior and physiology. Moreover, understanding the mechanisms of these sensory perceptions can be useful for developing chemical signal-based strategies that improve the integrated management of this species.

## AUTHORS CONTRIBUTION

S.R.R, I.O.L., M.V.T., P.S.Q. designed the research., S.R.R, I.O.L., M.V.T., A.S., P.S.Q. performed the work, analysed and discussed the results and wrote the paper.

## COMPLIANCE OF ETHICAL STANDARDS

### Ethical approval

All the animals employed in this study dead by natural causes.

### Informed consent

No human subject was used in this study.

## ACKNOWLEDGEMENTS

The authors wish to thank the “Dirección Xeral de Gandaría, Agricultura e Industrias Agroalimentarias of the Consellería do Medio Rural of the XUNTA de GALICIA” for the financial, logistical support, and the trust placed in this project.

